# Dentate gyrus network regulation by somatostatin- and parvalbumin-expressing interneurons differentially impacts hippocampal spatial memory processing

**DOI:** 10.1101/2025.07.23.666335

**Authors:** Frank Raven, Anna A. Vankampen, Annie He, Sara J. Aton

**Affiliations:** Department of Molecular, Cellular, and Developmental Biology, University of Michigan, Ann Arbor, MI 48019

## Abstract

GABAergic interneurons regulate circuit dynamics in hippocampal structures such as CA1 that appear to be essential for memory processing. The dentate gyrus (DG) is known to play a role in pattern recognition and spatial working memory. However, the role of the DG in different stages of long-term spatial memory is poorly understood. Moreover, the roles of the predominant interneuron subtypes within the DG - somatostatin-expressing (SST+) and parvalbumin-expressing (PV+) - in different stages of memory processing are unknown. We tested how chemogenetic manipulation of DG SST+ and PV+ interneurons in mice influences the encoding, consolidation, and retrieval of hippocampus-dependent object-location memory (OLM). We find that activation of DG SST+ interneurons impairs both OLM encoding and retrieval, dramatically suppresses DG granule cell cFos expression, and (in the case of encoding) suppresses downstream CA1 network activity. Among individual mice, the degree of DG granule cell suppression is proportional to the extent of SST+ interneuron activation, and predicts the extent of OLM deficits. In striking contrast, PV+ interneuron activation selectively disrupts encoding, but not retrieval, of OLM, and minimally impacts DG or downstream hippocampal network activity. These findings demonstrate that regulation of the DG network by SST+ and PV+ interneurons differentially contributes to the various stages of spatial memory processing, and suggest that distinct network mechanisms are engaged in the hippocampus during each processing stage.

**Significance statement:** Neuronal activity within the dentate gyrus (DG) of the hippocampus is regulated by multiple populations of inhibitory interneurons. To understand how inhibitory regulation contributes to spatial memory processing, we experimentally activated two major classes of DG inhibitory interneurons in mice - either during spatial learning, spatial memory storage, or memory recall. We find spatial learning is disrupted by activation of either SST+ or PV+ interneuron populations, although the two manipulations differentially affect hippocampal activity. Somewhat surprisingly, activation of neither inhibitory population affects spatial memory storage, only SST+ interneuron activation disrupts recall, and the hippocampal activity patterns affected by inhibition differ between learning and recall. These data provide a clearer understanding of the circuits engaged by different steps of spatial memory processing.

## Introduction

GABAergic interneurons provide the main source of inhibition to principal neurons in the hippocampus, which are widely believed to serve as “engram” populations encoding new episodic and spatial memories ^1^. These interneurons can be classified into multiple subpopulations based on morphological, electrophysiological, and biomarker expression. In the hippocampus, the most common interneuron types include somatostatin-expressing (SST+) and parvalbumin-expressing (PV+) interneurons, which provide local inhibition to neighboring principal cells at the level of dendrites and somata, respectively ^2,3^. Available experimental and computational modeling data suggest that the precise balance of activity between excitatory principal neurons and these inhibitory populations (i.e., network excitatory/inhibitory [E/I] balance) is tightly regulated in the context of learning and memory ^4–6^. Both hippocampal SST+ and PV+ interneuron populations undergo synaptic structural plasticity in response to memory encoding, which appears to — at least transiently — alter this balance ^7,8^.

Recent data suggest that hippocampal PV+ and SST+ interneurons play an active role in contextual memory encoding. For example, PV+ interneurons in both CA3 and DG selectively reduce their activity during exploration of novel, relative to familiar, environments. At the same time, DG SST+ interneurons increase their activity, while those in CA1 and CA3 become less active during exploration of novel environments ^9^. DG PV+ interneurons are essential for granule cell place field remapping, and subpopulations of PV+ and SST+ interneurons throughout the hippocampal circuit have their own place fields ^9^. These findings suggest interneuron roles in spatial and contextual memory encoding, through inhibition or disinhibition of specific principal neurons. In support of this idea, hippocampal inputs from the brainstem and septum can selectively activate CA1 and DG SST+ interneurons during contextual fear conditioning (CFC) ^10,11^. Widespread suppression of CA1 SST+ (but not PV+) interneurons’ activity is sufficient to disrupt contextual fear memory (CFM) acquisition ^12^. In DG, SST+ interneurons’ activity gates the activity of DG granule cells during both CFM encoding and retrieval ^13^. Inhibition of subpopulations of DG SST+ interneurons suffices to both encode CFM context and support retrieval ^14^. These findings suggest that SST+ and PV+ populations form an essential component of the hippocampal engram ^1^.

Available data also suggest a role for interneurons in the regulation of hippocampal memory consolidation. In CA1, PV+ interneurons generate oscillatory network activity patterns essential for CFM consolidation ^15–17^; chemogenetic or optogenetic inhibition of these interneurons in the hours following CFC is sufficient to disrupt CFM storage. CFM consolidation is disrupted by chemogenetic activation of DG SST+ interneurons in the hours immediately following CFC, while chemogenetic inhibition of this population enhances memory ^18^. These effects are likely linked to the role of DG SST+ interneurons in gating activity among neighboring granule cells. The inhibited granule cell population includes context-encoding engram neurons, which are selectively reactivated during CFM consolidation ^19^. Spatial transcriptomic data from the hippocampus also indicates that in the first 6 h following CFC, mRNAs encoding excitatory synapse components are upregulated, while inhibitory synapse components are downregulated ^19^. Recent computational modeling studies ^4^ suggest that dynamic changes in E/I balance across post-learning non-REM and REM sleep are essential for strengthening memory traces, and keeping engram populations encoding different memories separated, during memory consolidation.

Within the hippocampal DG, both interneurons ^14^ and granule cells ^19–21^ play an important role in encoding environmental context during CFM processing. However, less is known about how E/I balance in the DG (or elsewhere in the hippocampus) regulates the processing of other types of memory, such as spatial memory. Because lesion studies suggest that the DG plays an essential role in pattern separation ^9^ and spatial working memory ^22^, we tested how manipulation of E/I balance within DG affects long-term object-location memory (OLM) encoding, consolidation, and retrieval. We find that chemogenetic activation of DG SST+ or PV+ populations differentially affects these stages of OLM processing, as well as hippocampal network activity patterns.

## Materials and Methods

### Animals and housing

3-5 month old transgenic male and female mice (*SST-CRE* [B6N.Cg-Sst^tm2.1^(SST–cre)^Zjh^; Jackson] or *PV-CRE* [B6;129P2-Pvalb^tm1^(cre)^Arbr^/J; Jackson]) were used for all experiments (n = 9 animals/mouse line/sex). All mice were maintained on a 12 h light/12 h dark cycle (lights on 9AM – 9PM) and under constant temperature (22 °C ± 2 °C). Throughout experiments, mice were group housed in standard cages with transparent filtering lids with standard paper bedding and beneficial enrichment (nestlets and enviropack nesting material). Food and water were available *ad libitum*. Mice were housed and tested in the same room. All animal husbandry and experimental procedures were approved by the University of Michigan Institutional Animal Care and Use Committee (IACUC, PRO00011982)

### General experimental design

*SST-CRE* and *PV-CRE* mice were divided into two groups, with each group containing 4-5 male and 4-5 female mice. One group underwent bilateral transduction of the dorsal DG as previously described ^18^ (see below) to express the designer receptor exclusively activated by designer drugs (DREADD) hM3Dq-mCitrine, in order to selectively activate either SST+ or PV+ interneurons during different memory processing stages. Control mice were similarly transduced to express EGFP. Four weeks following AAV transduction, mice were habituated to daily handling (5 days, 2 min/day) and underwent initial OLM training and testing. Four OLM trials were performed at one-week intervals, using different objects and locations as described previously ^23^, in order to test each stage of memory processing separately. An overview of the experimental design is represented in Figure 1A. The day prior to OLM training, mice were individually placed in the OLM arena for five min with no objects present, to familiarize themselves with (and habituate to) the OLM training environment. In each experiment, the OLM paradigm consisted of a learning/encoding trial (T1) and a testing/retrieval trial (T2) 24 h later. To test the effects of interneuron activation on memory encoding, DREADD agonist compound 21 (C21; Tocris; 3 mg/kg in 10% DMSO in saline) was injected i.p. 30 min prior to OLM training (optimizing brain and plasma levels during encoding; ^24,25^(Figure 1B). To test effects on memory consolidation, C21 was injected i.p. immediately following OLM training. To assess effects on memory retrieval, C21 was injected i.p. 30 min prior to OLM testing. Vehicle was injected i.p. 30 min prior to OLM training in all mice as a control experiment. At the conclusion of these experiments, mice underwent a final OLM training and were euthanized 90 min later, to quantify cFos expression in the hippocampus during OLM encoding. An additional naive cohort of hM3Dq and control adult animals (n = 6/group) was sacrificed 90 min after OLM retrieval, in order to characterize cFos expression patterns in the hippocampus during retrieval.

**Figure 1:**
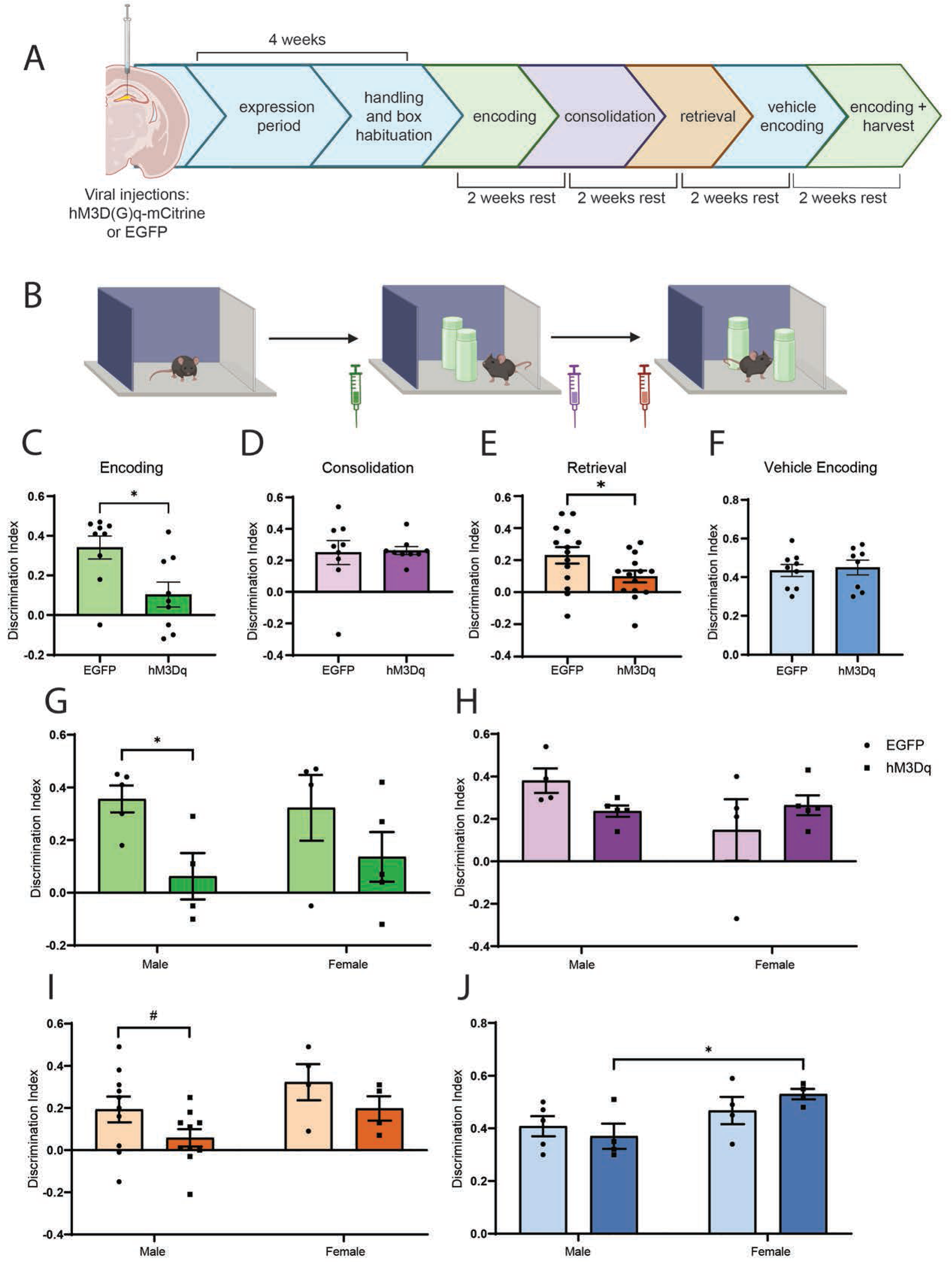
Effects of SST+ interneuron activation across different stages of spatial memory processing. (A) Male and female *SST-CRE* mice (n = 9 animals/group for encoding and consolidation; n = 14-15 animals/group for retrieval) were transduced to express Cre-dependent hM3Dq-mCitrine or EGFP (control) in DG. Beginning four weeks after viral transduction, SST+ interneurons were activated separately either during OLM encoding, consolidation, or retrieval, with two weeks of rest between OLM trials, which ended with a final trial in which vehicle was administered prior to encoding. (B) Timing of Compound 21 (C21) administration relative to OLMencoding, consolidation, or retrieval. Mouse interaction with the moved vs. unmoved object was quantified to generate the OLM discrimination index (DI). (C-E) SST+ interneuron activation significantly impaired OLM encoding (p = 0.013, Student’s t test) and retrieval (p = 0.046), but had no effect on consolidation (p = 0.881). (F) Vehicle administration did not affect OLM encoding (p = 0.755, n = 8-9 animals/group). (G-J) Sex-specific comparisons of OLM performance in EGFP-and hM3Dq-mCitrine-injected animals. (G) SST+ interneuron activation impaired OLM encoding in male mice only (p = 0.038, n = 4-5 animals/group; two-way ANOVA with AAV and sex as factors (2 way ANOVA, AAV×sex interaction: p = 0.564; main effect of AAV: p = 0.019; main effect of sex: p = 0.829). (H) Consolidation was unaffected across both sexes (n = 4-5 animals/sex/AAV; 2 way ANOVA, AAV×sex interaction: p = 0.105; main effect of AAV: p = 0.195; main effect of sex: p = 0.857). (I) A trend for impaired retrieval was apparent after SST+ interneuron activation for males only (p = 0.072, n = 10 males and 4 females/AAV; 2 way ANOVA, AAV×sex interaction: p = 0.957; main effect of AAV: p = 0.065; main effect of sex: p = 0.057). (J) Among vehicle-treated mice, OLM performance was significantly higher in hM3Dq-expressing females compared to hM3Dq-expressing males (p = 0.019, n = 4-5 animals/sex/AAV; 2 way ANOVA, AAV×sex interaction: p = 0.245; main effect of AAV: p = 0.772; main effect of sex: p = 0.020). For all panels, #p < 0.1, *p < 0.05.

### Viral transduction surgeries

Adeno-associated viruses (AAVs) were used to express either the depolarizing DREADD hM3dq-mCitrine (pAAV-hSyn-DIO-hM3D(Gq)-mCitrine; Addgene #50454-AAV8) or EGFP (pAAV-hSyn-DIO-EGFP; Addgene #50457-AAV8) in neurons in a CRE-dependent manner. Stereotaxic surgeries targeting vectors to DG were performed as described previously ^18^. Briefly, *SST-CRE* and *PV-CRE* mice were anesthetized with isoflurane, the skull overlying dorsal hippocampus was exposed, and holes were drilled bilaterally to introduce a hamilton syringe needle into DG for stereotaxic AAV delivery at the following coordinates A/P -2.0 mm, M/L ±1.5 mm, D/V -2.0 mm. A total volume of 1 µl (titer ≥ 1×10¹³ vg/mL) was injected per hemisphere at a rate of 100 nl/min. Following injections, the syringe needle was left in place for 2 min before being gradually removed, and incision margins were closed using nylon sutures and tissue adhesive (Vetbond; 3M). Each mouse was administered ketofen (5 mg/kg, s.c.) prior to surgery, and 24 and 48 hours post-surgery.

### The object-location memory paradigm

The OLM behavioral paradigm relies on mice’s innate preference for investigating objects that have been moved from their prior location. Measures of this bias for spatial novelty are frequently used to assess hippocampus-dependent spatial memory ^23,26,27^. The rectangular OLM arena was a square 40 cm x 30 cm x 30 cm (length x width x height) made from PVC, with grey walls, an open top, and a transparent bottom. Two walls of the arena had a visual spatial cue (e.g., a vertical line pattern) to facilitate spatial orientation within the arena. Different sets of objects were used that varied in texture and color (see Supplementary Figure1). Two identical glass or plastic objects were randomly assigned to each mouse; a different object set was used for each OLM experiment. At lights on (ZT0), 24 h prior to OLM training, each mouse was placed in the OLM arena and was allowed to explore the empty box without objects for 5 min. During OLM training at ZT0 the following day, each mouse was allowed to freely explore a pair of identical objects placed symmetrically within the arena for 10 min. OLM testing occurred 24 h after training, at ZT0. During testing, each mouse was returned to the arena containing the two objects, with one of the two objects displaced from its initial location, and was allowed to freely explore the arena and objects for 10 min. Between each of the training and testing trials, the objects and object box were cleaned using 10% ethanol. Exploratory behavior during training and testing was recorded using a video camera mounted directly above the arena. Briefly, exploration was defined as directing the nose toward the object while being positioned within a 2 cm radius of the object. This measurement included time spent touching the object, but excluded time spent climbing on the object. During the test, the time spent exploring the stationary and moved objects are measured. We calculated the discrimination index (DI) by subtracting the exploration time of the stationary object from that of the moved object, then dividing the result by the total exploration time of both objects. Hence, the DI is calculated by: (moved object exploration time - stationary object exploration time)/(moved object exploration time + stationary object exploration time). These values were scored in a semi-automated manner from trial videos using Ethovision XT 16 (Noldus Information Technology, The Netherlands). Object exploration times were independently verified by manual scoring of two blinded experimenters.

### Tissue processing and quantitative immunohistochemistry

To collect hippocampal tissue in the context of OLM processing, mice were administered euthasol euthanasia (pentobarbital sodium and phenytoin sodium, Virbac) and were transcardially perfused with phosphate-buffered saline (PBS) followed by 4% paraformaldehyde (PFA) in PBS. In the first set of experiments, brain samples were collected either 90 min after the OLM training (i.e., encoding), or 90 min after OLM testing (i.e., retrieval). Following perfusion, the brains were post-fixed in 4% PFA for 24 hours at 4⁰C, then transferred to PBS. Brains were sectioned coronally at 80 µm thickness using a Leica vibratome.

Free-floating brain sections containing dorsal hippocampus were blocked overnight at 4⁰C in a PBS solution containing 5% normalized goat serum (NGS) and 1% Triton X-100, then were incubated for 72 hours at 4⁰C in PBS with 5% NGS, 0.5% Triton X-100, and primary antibodies. Sections were rinsed three times at room temperature (1 h each) in PBS containing 1% NGS and 0.5% Triton X-100, followed by incubation for 72 hours at 4⁰C in PBS with 5% NGS, 0.5% Triton X-100, and secondary antibodies. Following an additional three 1-h rinses at room temperature in PBS containing 1% NGS and 0.5% Triton X-100, sections were mounted in PBS and coverslipped using Prolong Diamond mountant (Thermo Fisher).

For immunohistochemistry on samples from *SST-CRE* animals, chicken anti-EGFP (1:750, Rockland; AB_1537403), mouse anti-PV (1:500, Chemicon, AB_477329), and rabbit anti-cFos (1:500, Cell Signaling Technology; AB_2247211) were used as primary antibodies, and goat anti-chicken Alexa488 (1:1000, Invitrogen, AB_2534096), goat anti-mouse Alexa594 (1:500, Invitrogen, AB_2534091), and goat anti-rabbit Alexa647 (1:500, Invitrogen, AB_2535813) were used as secondary antibodies. For samples from *PV-CRE* animals, chicken anti-EGFP (1:750, Rockland, AB_1537403) and rabbit anti-cFos (1:500, Cell Signaling Technology, AB_2247211) were used as primary antibodies, and goat anti-chicken Alexa488 (1:1000, Invitrogen, AB_2534096) and goat anti-rabbit Alexa594 (1:500, Invitrogen, RRID:AB_2534095) were used as secondary antibodies, along with lectin Wisteria floribunda Lectin-Cy5 (WFA/WFL; 1:750, BioWorld, #21761157).

Images were acquired from immunolabeled sections using an SP8 confocal immunofluorescence microscope (Leica). Images of the DG hilus, CA1, and CA3 subregions were imaged separately at 10x. Acquisitions spanned 32 µm in depth (Z-step size of 8 µm) with identical exposure times for each specific immunostaining between and within samples. Quantification of immunolabeling was carried out in a blinded, semi-automated manner using ImageJ software. Area for each subregion was used to calculate the density of cell expression of somatostatin, parvalbumin, cFos, or WFA, as well as colocalization. ImageJ was also used to calculate the percentage SST+ or PV+ populations positively immunolabeled with cFos or WFA.

### Statistical analyses

Experimenters were blinded to treatment conditions while performing data analyses. Statistical analyses were carried out using GraphPad Prism10 (v10.2.3). In comparisons between EGFP- and hM3Dq-mCitrine-injected animals, we used a two-sample t-test. In comparisons between male and female mice expressing EGFP or hM3Dq, we used a two-way ANOVA to calculate statistical differences, followed by a Tukey test for multiple comparisons, with sex and AAV as independent variables. Trends were reported when p < 0.1, and statistically significant differences were reported when p < 0.05. All data are presented as mean ± S.E.M.

## Results

### SST+ interneuron activation impairs spatial memory encoding and retrieval, but not consolidation

To test how selective DG SST+ interneuron activation affects OLM encoding, consolidation, and retrieval, we carried out four separate OLM trials (each using different objects and object placement patterns) to target each stage individually ^23^. C21 was administered to *SST-CRE* mice expressing either the depolarizing DREADD hM3Dq or EGFP at different stages during each trial: either 30 min before OLM training (encoding), immediately after OLM training (consolidation), or 30 min before OLM testing (retrieval), as shown in Figure 1A-B. For each mouse, in the fourth OLM trial, vehicle (10% DMSO in saline) was administered 30 min before OLM training. A discrimination index (DI) was calculated from each testing trial as an indicator of spatial memory performance, where a higher DI score reflects preferential interaction with the object that had been moved from its prior location (i.e., a stronger spatial memory).

Using DI as our memory index, we found that SST+ interneuron activation during OLM training (i.e., encoding) significantly impaired OLM performance, affecting male mice more severely than females (Figure 1C, G). This encoding effect was specific to hM3Dq mice administered C21; mice administered a vehicle prior to training successfully encoded OLM (Figure 1F). In contrast, C21 administration to activate SST+ interneurons immediately following training did not affect OLM consolidation, in either males or females (Figure 1D, H). SST+ interneuron activation just prior to testing significantly impaired OLM retrieval (Figure 1E, I), with reductions in DI that were more pronounced in male mice. Interestingly, we found a trend for a main effect of sex, where female mice performed better than males overall (Figure 1E, I). Again, female mice showed a higher DI, indicating better memory performance compared to males (Figure 1J). These data suggest that selective activation of SST+ interneurons in the DG impairs encoding and retrieval, but not the consolidation, of spatial memory.

### SST+ interneuron activation alters DG activity patterns associated with spatial memory encoding and retrieval

Our recent data indicate that hippocampal principal neuron activation patterns associated with successful contextual fear memory consolidation and retrieval are subregion specific ^19,28,29^. To clarify how SST+ interneuron activation affects the activity of principal neurons in the context of hippocampal OLM processing, we quantified cFos protein expression as an indicator of neuronal activity during memory encoding and retrieval. C21 was administered to *SST-CRE* mice (expressing hM3Dq or EGFP) either 30 min before OLM training or 30 min prior to OLM retrieval, and brain tissue was collected 90 min post-training or post-retrieval for immunostaining. Immunohistochemistry was used to visualize and quantify SST+ interneurons, PV+ interneurons, and cFos expression within the DG hilus, superior blade, and inferior blade (Figure 2A), as well as areas CA1 and CA3 (Figure 3A, 4A). In the DG, immunostaining confirmed successful chemogenetic activation of hilar SST+ interneurons during OLM encoding, with more interneurons co-expressing cFos in hM3Dq-expressing mice compared to EGFP control mice (Figure 2B-C). SST+ interneuron activation decreased cFos expression among granule cells in both the DG superior and inferior blade, with greater suppression of activity in the inferior blade (Figure 2D). This effect differed between male and female mice, with reductions in inferior blade cFos expression being more pronounced in females (Figure 2E). cFos expression among DG granule cells did not differ between male and female EGFP expressing mice, which had no SST+ interneuron activation nor encoding impairment (Figure 2E). No significant effects on cFos expression among DG PV+ interneurons were observed (Figure 2F).

**Figure 2:**
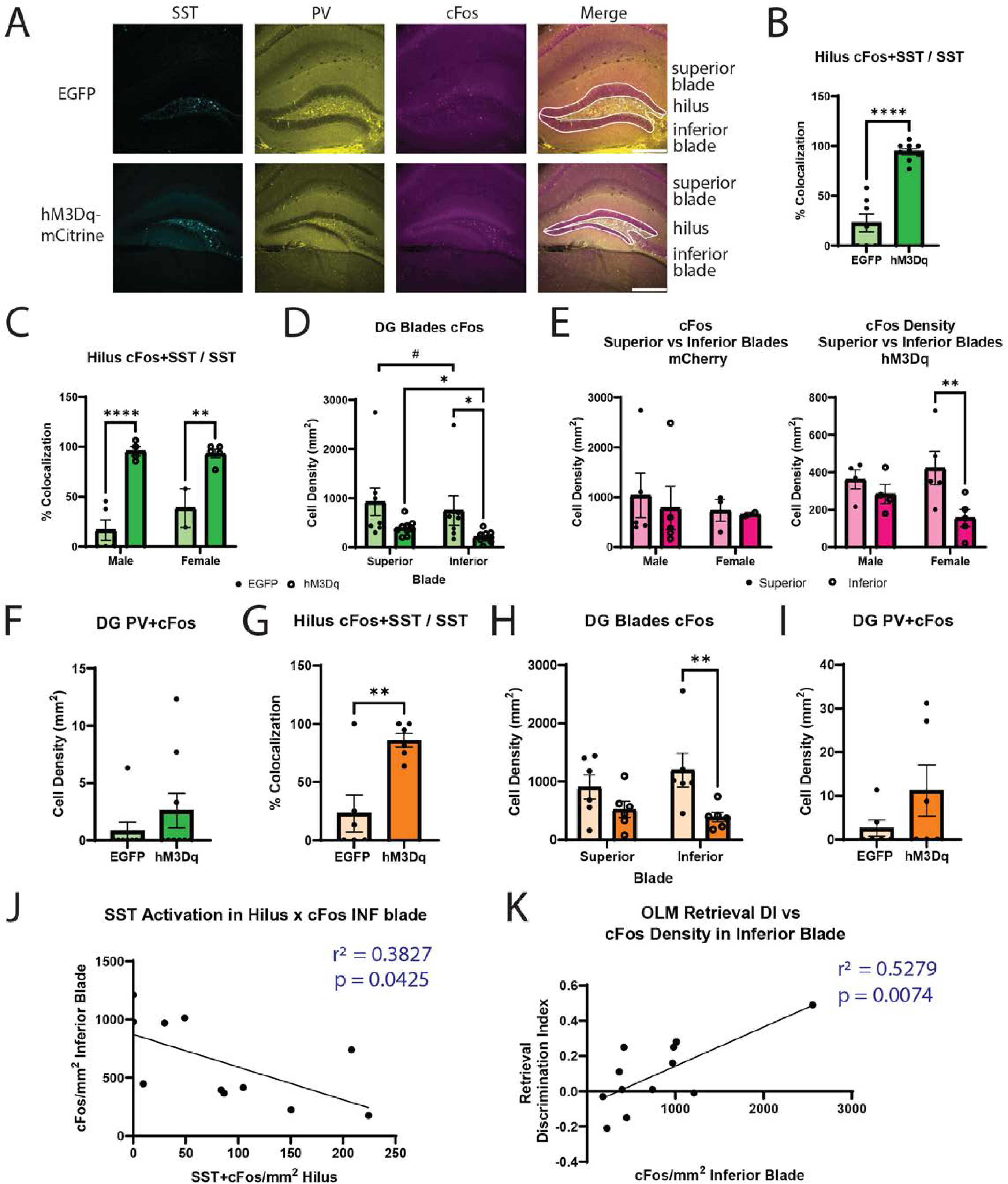
Activation of DG SST+ interneurons during spatial memory encoding vs. retrieval differentially alters DG network activity. Quantification of cFos expression in the DG associated with OLM encoding (B-H; n=18) or retrieval (I-N; n=12). (A) Max projection of DG in EGFP- and hM3Dq-mCitrine expressing *SST-CRE* mice following immunohistochemistry to visualize virally-transduced SST+ interneurons (cyan), PV+ interneurons (yellow), and cFos (magenta). Scale bar = 300μm. (B) Chemogenetically-activated SST+ interneurons expressed significantly higher cFos during encoding (****p < 0.0001, n = 7-9 animals/group). (C) These effects were present in both males and females (males: ****p < 0.0001, n = 2-5 animals/group; females: **p = 0.002, n = 2-5 animals/group; 2 way ANOVA, AAV×sex interaction: p = 0.203; main effect of AAV: p < 0.0001; main effect of sex: p = 0.309). (D) Activation of SST+ interneurons during encoding significantly reduced cFos expression among DG granule cells in the inferior blade, while encoding-driven cFos expression was lower in the inferior vs. superior blade across both groups (2-way ANOVA, 2 way ANOVA, AAV×blade interaction: p = 0.871; main effect of AAV: p = 0.053; main effect of blade: p = 0.006, n = 7-9/group). *p < 0.05, ^#^p = 0.058. (E) This effect of SST+ interneuron activation on granule cell cFos expression was stronger in female mice (*p = 0.0043, n = 4-5/group) than male mice (p = 0.3138, n = 4-5/group). (F) SST+ interneuron activation during encoding did not significantly affect cFos expression among PV+ interneurons (p = 0.321, n = 8-9/group). (G) Activation of SST+ interneurons during retrieval activated SST+ interneurons in the DG hilus (**p = 0.004, n = 6/group). (H) Retrieval-associated SST+ interneuron activation suppressed cFos expression among DG granule cells in inferior blade (p = 0.009, n = 6/group), with more modest effects on the superior blade (p = 0.183, n = 6/group), DG granule cells. (I) SST+ interneuron activation did not significantly alter DG PV+ interneuron cFos expression associated with retrieval, although there was a trend for increased expression (p = 0.193, n = 6/group). (J) Density of cFos-expressing SST+ interneurons is inversely correlated with cFos expression in the inferior blade granule cells (p = 0.043). (K) cFos expression among inferior blade granule cells was positively correlated with OLM discrimination index during retrieval (p = 0.007). Two-group comparisons were analyzed using unpaired t-tests, while four-group designs were assessed using two-way ANOVA with AAV and sex as independent factors.

We next tested whether SST+ interneuron activation during OLM retrieval similarly affected the activity pattern in DG. Chemogenetic activation of hilar SST+ interneurons was again confirmed based on the higher number of neurons co-expressing SST and cFos in hM3Dq mice compared to EGFP controls (Figure 2G). Similar to effects on network activity during encoding, chemogenetic SST+ interneuron activation during retrieval significantly reduced granule cell cFos density, with strongest effects within the inferior blade (Figure 2H), and no effects on cFos expression among PV+ interneurons (Figure 2I). We also found that there was an inverse relationship between cFos expression among SST+ interneurons and cFos expression among DG inferior blade granule cells during OLM retrieval (Figure 2J), but not cFos expression in superior blade granule cells. This supports the idea that these interneurons gate granule cell activity in the context of OLM processing, similar to their role in CFM encoding and retrieval ^13,18^, and that they preferentially inhibit inferior blade granule cells. We also found that individual animals’ OLM performance (DI) was predicted by retrieval-associated cFos in inferior blade granule cells, across both treatment groups (Figure 2K). Together, these data suggest that the DG inferior blade, which is a major target of SST+ interneuron-mediated inhibition, plays a vital role in OLM retrieval.

### DG SST+ interneuron activation alters CA3 and CA1 activity patterns associated with spatial memory encoding and retrieval

Hilar SST+ interneurons affect local circuits within the DG, the output of which affects neuron populations in hippocampal CA1 and CA3 ^30^. Because these structures also have important roles in memory processing, we further quantified images of CA1 (including the stratum oriens [SO] and stratum radiatum [SR]; Figure 3A) and CA3 (Figure 4A). SST+ interneuron activation reduced overall CA1 cFos expression during encoding (Figure 3B). This effect was spatially uniform, without clear differences between individual cell layers (Figure 3C). When separated by sex, this significant difference was only seen in the female group, and not in males (Figure 3D). SST+ interneuron activation during encoding caused no overall change in cFos expression among PV+ interneurons in CA1 (Figure 3E). However, the proportion of cFos+ PV+ interneurons was significantly lower in control females compared to males (Figure 3F). In contrast to the effects observed during encoding, SST+ interneuron activation during retrieval did not alter overall cFos expression in CA1 (Figure 3G). In the SR, there was a trend for decreased cFos expression, but no differences were observed in the SO (Figure 3H). No differences in PV+ interneuron cFos expression were observed between groups during retrieval (Figure 3I).

**Figure 3:**
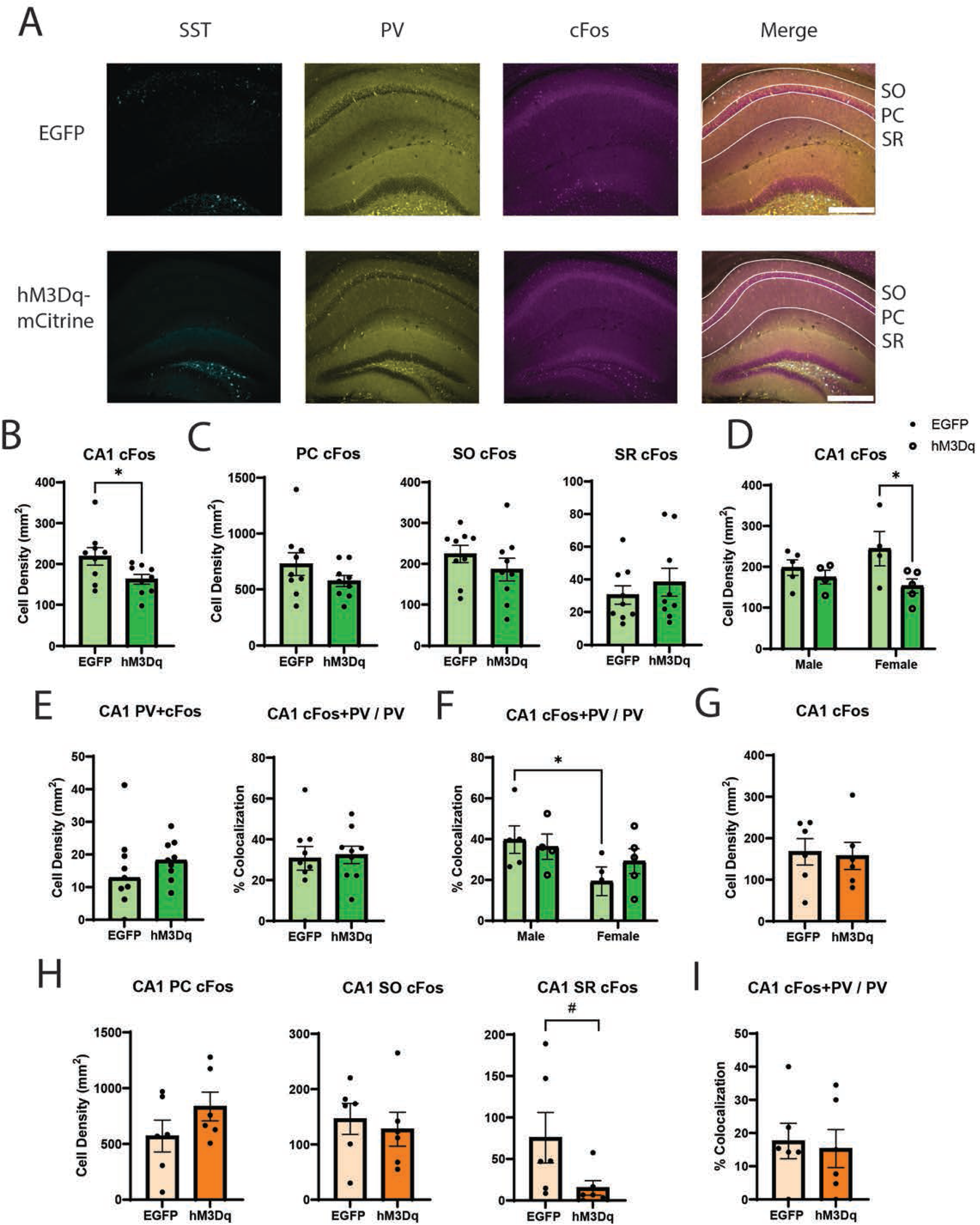
Activation of DG SST+ interneurons during spatial memory encoding vs. retrieval differentially alters CA1 network activity. Quantification of CA1 cFos expression during encoding (B-F; n=18) and retrieval (G-I; n=12) in EGFP- or hM3Dq-mCitrine-expressing *SST-CRE* mice. (A) Max projection images of immunolabeling for transduced SST+ interneurons in DG (cyan) and PV+ interneurons (yellow), and activity marker cFos (magenta) in CA1, following OLM encoding. For quantification, CA1 was subdivided into the pyramidal cell (PC) layer, stratum oriens (SO), and stratum radiatum (SR). Scale bar = 300 μm. (B-C) Chemogenetic activation of DG SST+ interneurons during encoding decreased overall CA1 cFos expression in CA1 (*p = 0.037, n = 9/group), although this effect was not significant when specific CA1 subregions were quantified separately (PC: p = 0.199, n = 9/group; SO: p = 0.292, n = 9/group; SR: p = 0.455, n = 9/group). (D) Effects of SST+ interneuron activation on encoding-associated CA1 cFos expression were present in female (*p = 0.020, n = 4-5/group), but not male (p = 0.514, n = 4-5/group) mice (2 way ANOVA, AAV×sex interaction: p = 0.190; main effect of AAV: p = 0.036; main effect of sex: p = 0.620). (E-F) Chemogenetic activation did not alter CA1 PV+ interneuron cFos expression during encoding (E: left: p = 0.494, n = 9/group; right: p = 0.820, n = 9/group), although PV+ interneuron activity differed between male and female control mice (F: *p = 0.046, n = 4-5/group; 2 way ANOVA, AAV×sex interaction: p = 0.425; main effect of AAV: p = 0.346; main effect of sex: p = 0.064). (G) When DG SST+ interneurons were activated during OLM retrieval, cFos expression in CA1 was not affected (p = 0.8331, n = 6/group). (H) SST+ interneuron activation during retrieval did not alter cFos expression in the PC (p = 0.1978, n = 6/group) or SO (p = 0.6641, n = 6/group), but trended toward reducing expression in the SR (^#^p = 0.085, n = 6/group). (I) Chemogenetic activation during retrieval did not alter PV+ interneuron cFos expression in CA1 (p = 0.776, n = 6/group). Two-group comparisons were analyzed using unpaired t-tests, while four-group designs were assessed using two-way ANOVA with AAV and sex as independent factors.

In hippocampal area CA3 (Figure 4A), cFos expression associated with encoding did not change as a function of chemogenetic manipulation, and no sex differences were observed (Figure 4B). While chemogenetic activation of SST+ interneurons did not alter cFos expression among CA3 PV+ interneurons (Figure 4C), as was true in CA1, male control mice showed a higher level of PV+ interneuron cFos expression during encoding than female mice (p < 0.01; Figure 4D). In addition, strong trends for directionally opposite effects were observed between male and female mice. In male mice, SST+ interneuron activation during encoding tended to reduce PV+ interneuron cFos expression while in females, it tended to increase it (p < 0.1; Figure 4D). When SST+ interneurons were activated during retrieval, there were no significant changes in overall cFos expression or PV+ interneuron cFos expression (Figure 4E-F).

**Figure 4:**
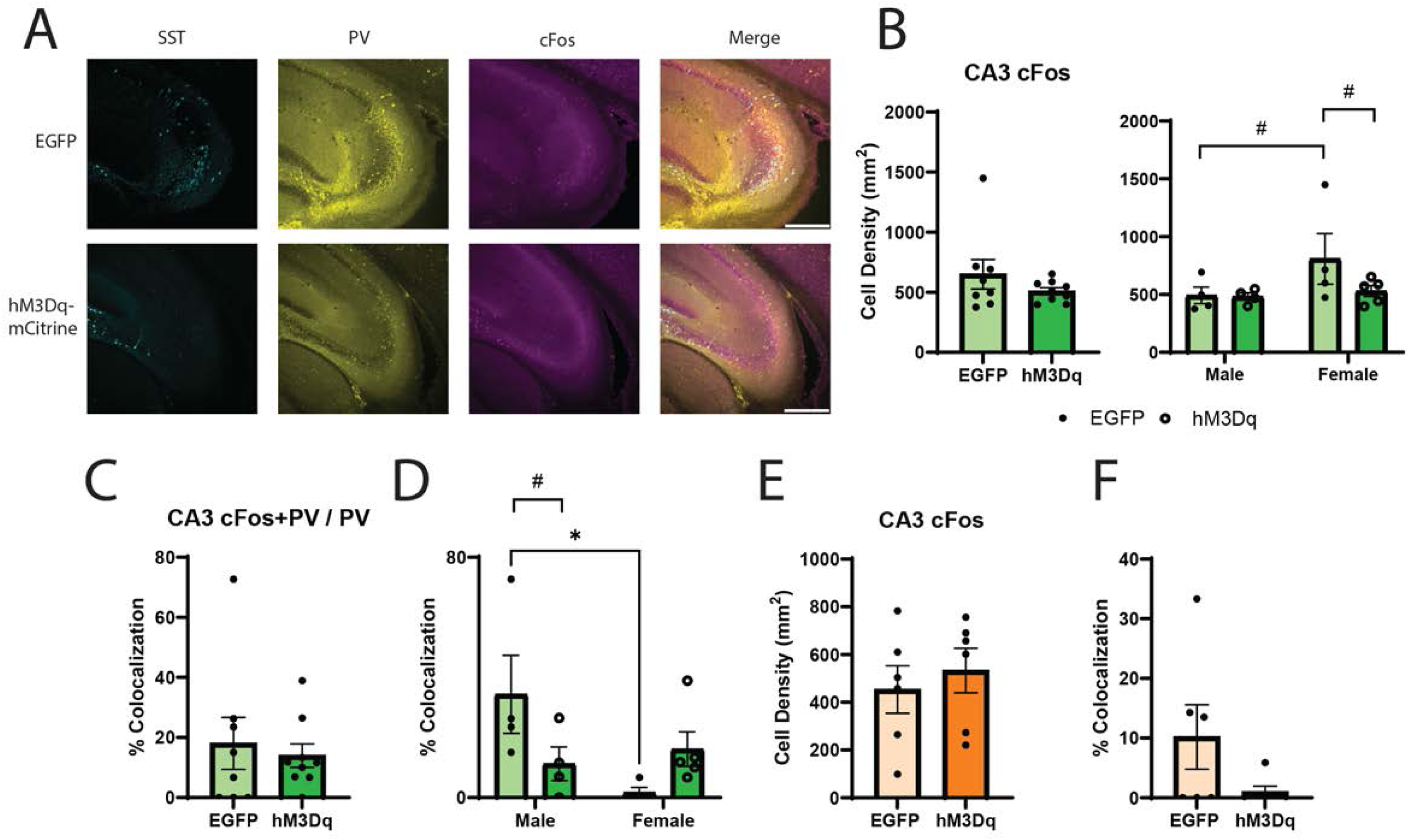
Activation of DG SST+ interneurons during encoding, but not retrieval, subtly alters CA3 network activity. Quantification of cFos expression in CA3 during OLM encoding (B-D; n=18) and retrieval (E-F; n=12) in EGFP- or hM3Dq-mCitrine-expressing *SST-CRE* mice. (A) CA3 max projection to visualize immunolabeling for transduced SST+ interneurons in DG (cyan), PV+ interneurons (yellow), and cFos (magenta) following encoding. Scale bar = 300 μm. (B) Chemogenetic SST+ interneuron activation during encoding did not alter total cFos expression in CA3 (p = 0.252, n = 8-9/group), although cFos tended to be reduced by activation in female mice only (male: p = 0.952, n = 4-5/group; female: p = 0.096, n = 4-5/group; 2 way ANOVA, AAV×sex interaction: p = 0.255; main effect of AAV: p = 0.222; main effect of sex: p = 0.129). CA3 cFos expression in female control mice trended higher than in male control mice (p = 0.075, n = 4-5/group). (C-D) SST+ interneuron activation during encoding did not affect PV+ interneuron cFos expression in CA3 across both sexes, (C: p = 0.662, n = 8-9/group). However, density of cFos+PV+ interneurons shows a decreasing trend in males (D: ^#^p = 0.054, n = 4-5/group), and an increasing trend in females (D: p = 0.187, n = 4-5/group). Among control (EGFP-expressing) mice, PV+ interneuron activation during encoding differed significantly between males and females (D: *p = 0.010, n = 4-5/group). 2 way ANOVA, AAV×sex interaction: p = 0.027; main effect of AAV: p = 0.574; main effect of sex: p = 0.088. (E-F) Chemogenetic SST+ interneuron activation during retrieval did not alter overall CA3 cFos expression (p = 0.5708, n = 6/group), nor the cFos expression of CA3 PV+ interneurons (left: p = 0.1111, n = 6/group; right: p = 0.1243, n = 6/group). Two-group comparisons were analyzed using unpaired t-tests, while four-group designs were assessed using two-way ANOVA with AAV and sex as independent factors.

### DG PV+ interneuron activation disrupts spatial memory encoding, but not consolidation or retrieval

In parallel with studies assessing effects of DG SST+ interneuron activation, we also tested the effects of DG PV+ interneuron activation on OLM encoding, consolidation, and retrieval. Using methods described above, *PV-CRE* transgenic mice underwent transduction to express depolarizing DREADD hM3Dq or EGFP in DG PV+ interneurons. Each mouse underwent four OLM trials as described above. As was observed with SST+ interneuron activation, chemogenetic PV+ interneuron activation during encoding significantly impaired OLM. However, in contrast to the trends for sex differences observed with SST+ interneuron activation, PV+ interneuron activation had a stronger effect in female mice than in males (Figure 5A-B). This encoding effect was specific to hM3Dq mice administered C21; mice administered a vehicle prior to training successfully encoded OLM (Figures 5E). PV+ interneuron activation, like SST+ interneuron activation, did not affect OLM consolidation, but unlike SST+ interneuron activation, PV+ interneuron activation had no effect on retrieval in either males or females (Figures 5C-D). These results suggest that DG PV+ interneurons specifically regulate the encoding phase of spatial memory processing.

**Figure 5:**
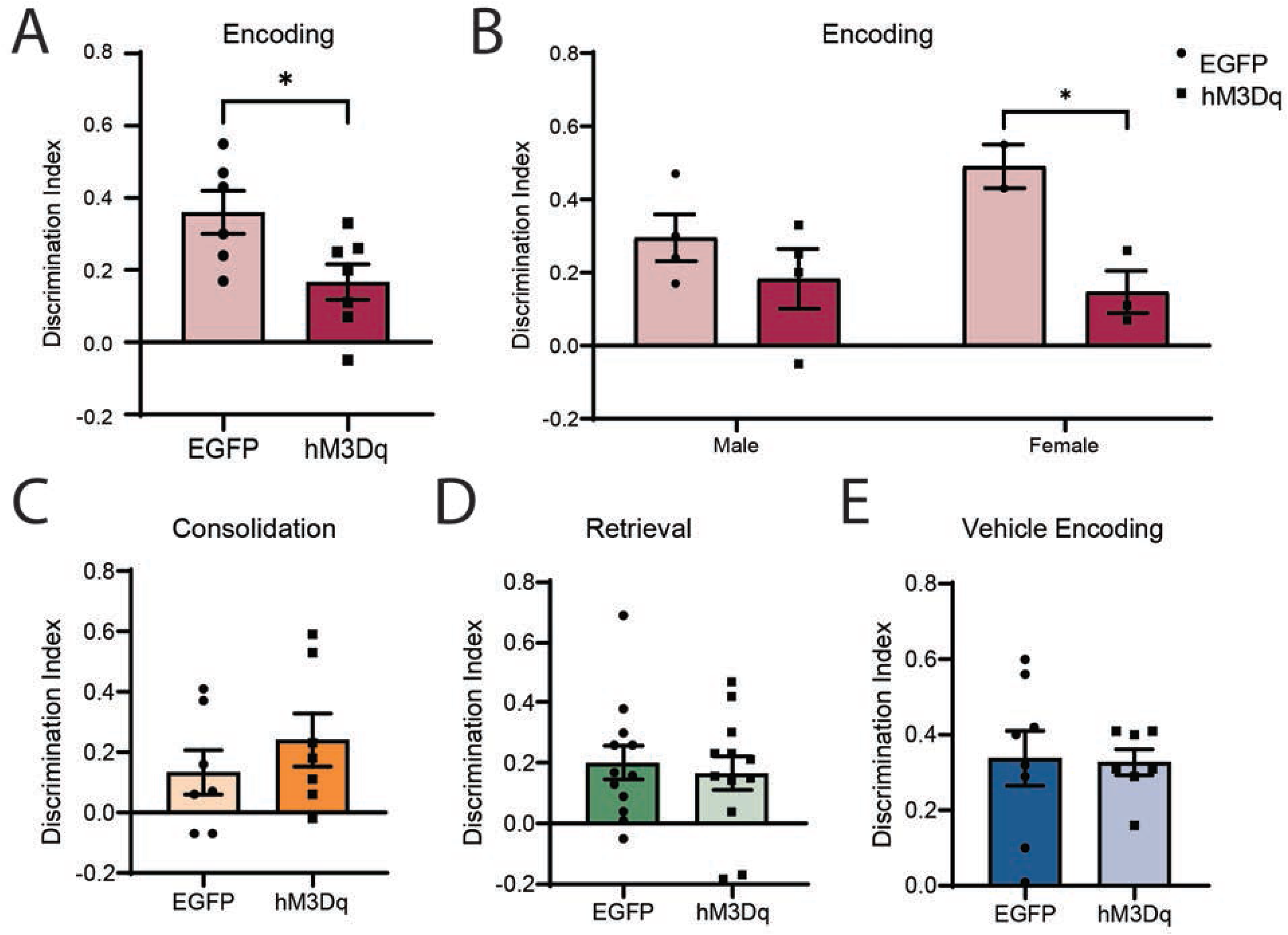
DG PV+ interneuron activation impairs encoding, but not consolidation or retrieval, of spatial memory. (A-C) Comparison of DIs in EGFP- and hM3Dq-mCitrine-expressing *PV-CRE* mice when PV+ interneurons were selectively activated during each phase of OLM processing. (A) PV+ interneuron activation during encoding significantly impaired spatial memory (p = 0.029, Student’s t-test, n = 6-7 animals/group), but had no effect on (B) consolidation (n = 7 animals/group) or (C) retrieval (n = 12 animals/group). (D) Vehicle administration had no effect on encoding (n = 7-8 animals/group). (E) PV+ interneuron activation effects on encoding were significant among female mice only (p = 0.019, Tukey test for multiple comparisons, n = 2-4 animals/sex/AAV).

### Activation of DG PV+ interneurons minimally affects hippocampal network activity patterns during OLM encoding and retrieval

To better understand differences in the behavioral effects of SST+ and PV+ interneuron activation, we quantified changes to hippocampal network activity when DG PV+ interneurons were activated during OLM encoding and OLM retrieval. C21 was administered to *PV-CRE* mice (expressing hM3Dq or EGFP) 30 min prior to OLM training, and brain tissue was collected 90 min after training for immunostaining for cFos and a marker of PV+ interneuron synaptic plasticity-regulating perineuronal nets (PNNs), wisteria floribunda lectin (WFA). Verifying the effects of the chemogenetic manipulation, we found that PV+ interneuron activation during encoding increased the percentage of active (cFos+) virus-labeled PV+ interneurons throughout the DG (Figure 6A-D). Chemogenetic activation also led to a trend toward an increased percentage of PV+ interneurons surrounded by PNNs (Figure 6E-G). However, unlike the effects of SST+ interneuron activation, chemogenetic PV+ interneuron activation did not affect the overall pattern of cFos expression in DG granule cells during encoding (Figure 6H-J). We also observed no differences in overall cFos expression, or cFos expression in interneurons with PNNs, in either CA1 (Figures 7A-C) or CA3 (Figures 7D-F). Taken together, these data suggest that activation of DG PV+ interneurons during memory encoding disrupts OLM without significantly altering encoding-related dorsal hippocampal network activity.

**Figure 6:**
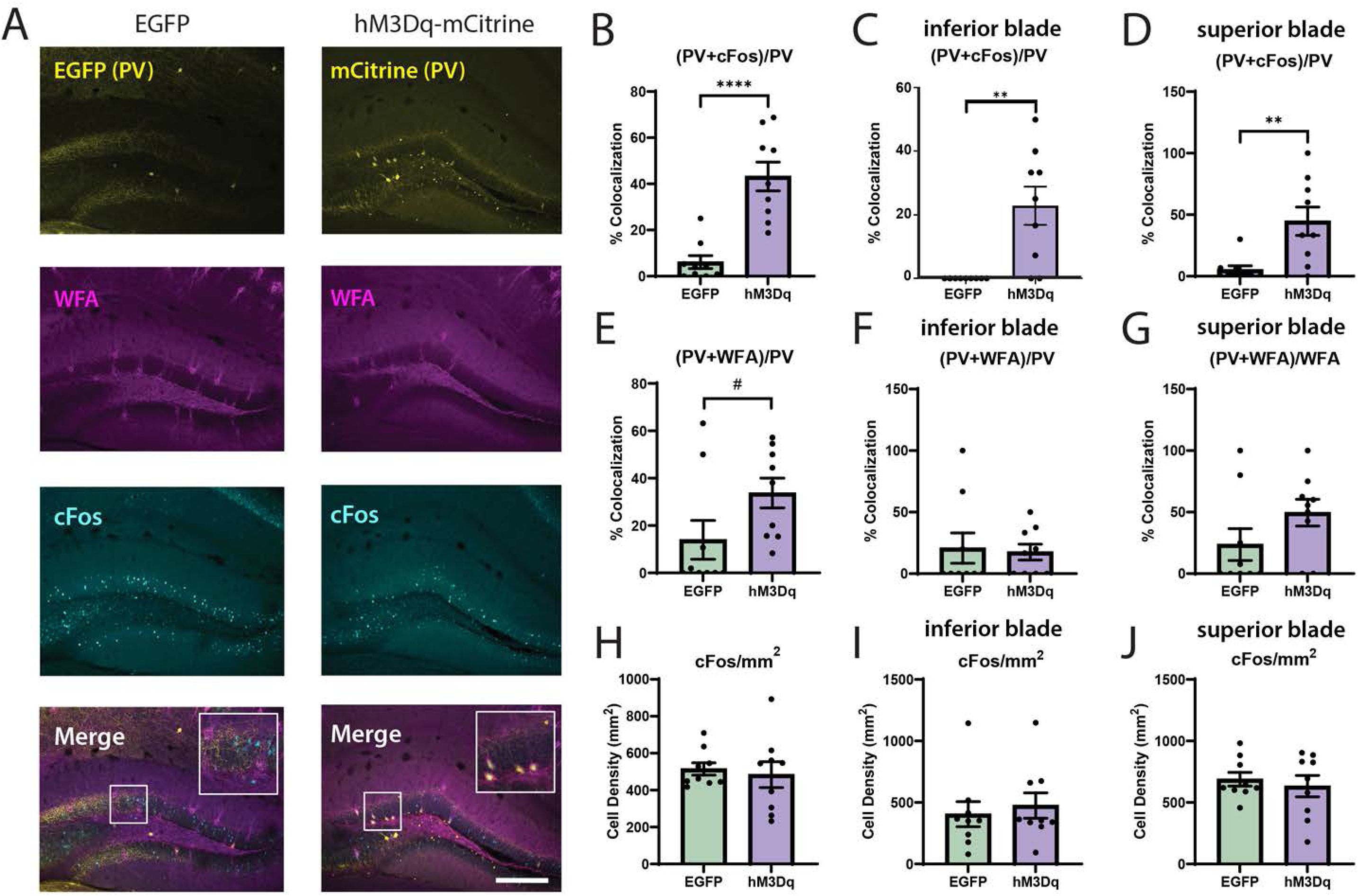
PV+ interneuron activation during OLM encoding increases DG PV+ interneuron perineuronal nets, but does not affect DG granule cell activity. (A) Representative images of DG in EGFP- and hM3Dq-mCitrine-expressing *PV-CRE* mice, showing immunolabeling for transduced PV+ interneurons in DG (yellow), perineuronal nets (Wisteria floribunda lectin [WFA]; magenta), and activity marker cFos (cyan) following OLM encoding. Scale bar = 250 µm. (B) PV+ interneuron activation increased their cFos expression during encoding (p < 0.0001, Student’s t-test, n = 9 mice/group), in both the inferior (C; p = 0.002) and superior blade of DG (D; p = 0.004). 2 way ANOVA, AAV×blade interaction: p = 0.163; main effect of AAV: p < 0.001; main effect of blade: p = 0.031. (E) The percentage of PV+ neurons co-labeled with WFA showed a trend towards an increase following chemogenetic activation (^#^p = 0.075), largely driven by changes in the superior blade compared to the inferior blade (F; p = 0.819; G; p = 0.285). 2 way ANOVA, AAV×blade interaction: p = 0.030; main effect of AAV: p = 0.612; main effect of blade: p = 0.019. (H) The density of cFos-expressing granule cells did not differ between groups (p = 0.946), although expression was generally higher during encoding in the superior blade (I-J). 2 way ANOVA, AAV×blade interaction: p = 0.455; main effect of AAV: p = 0.944; main effect of blade: p = 0.017).

**Figure 7:**
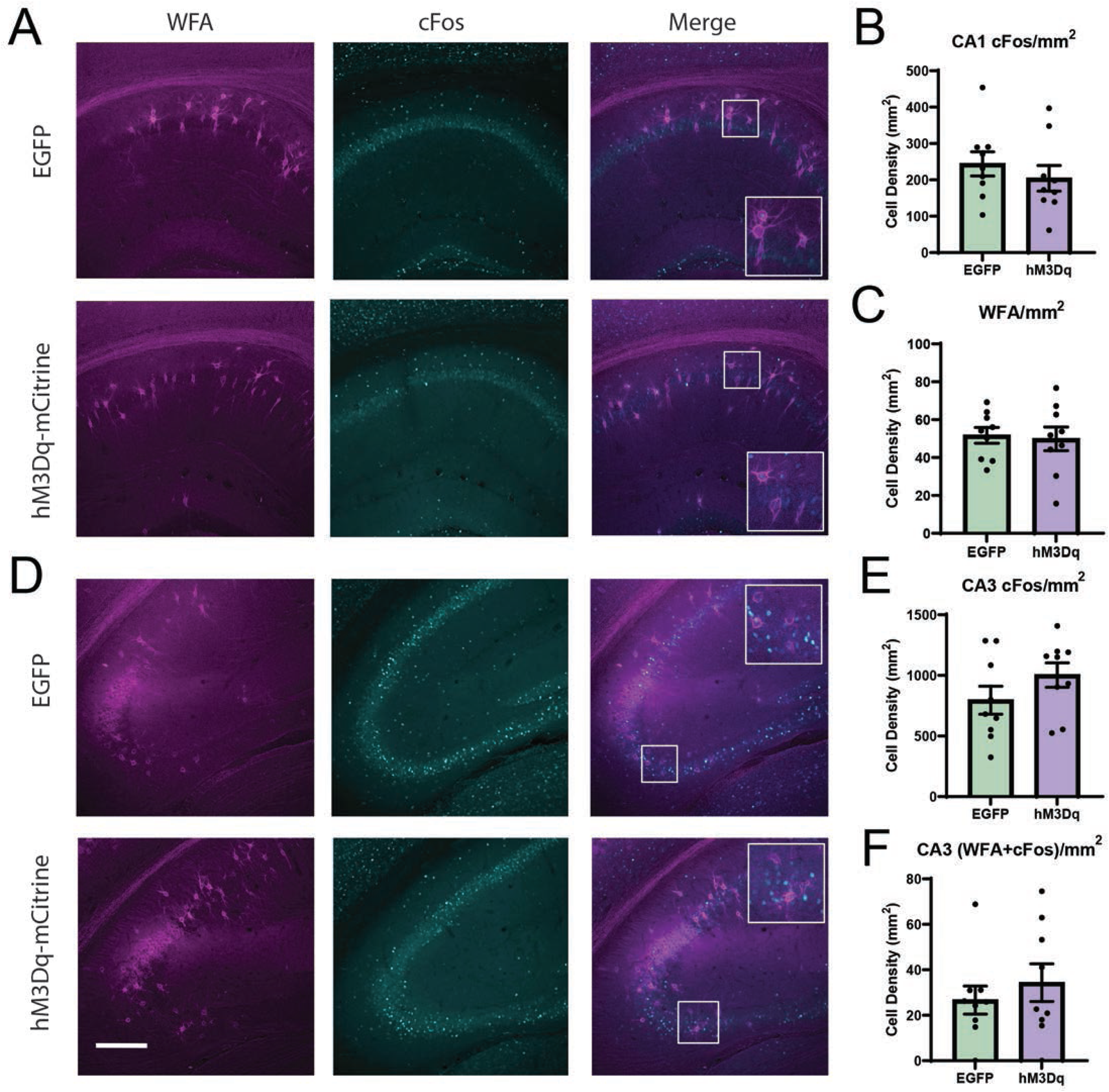
DG PV+ interneuron activation does not affect CA1 or CA3 network activity during OLM encoding. (A) Representative images of CA1 following encoding in EGFP- and hM3Dq-mCitrine-expressing *PV-CRE* mice, showing WFA (magenta) and cFos (cyan) immunolabeling. Scale bar = 250 µm. There were no differences in CA1 between groups in the density of cFos-positive cells (B; p = 0.423), or the density of WFA labeled cells across all subregions in CA1 (C; p = 0.812). (D) Representative images of CA3 WFA (magenta) and cFos (cyan) immunolabeling. Scale bar = 250 µm. There were no differences between groups for the density of cFos-positive cells (E) or the density of WFA/cFos co-labeled cells in CA3 (F). All statistical analyses were performed using the student’s t-test. n = 8-9 animals/group.

Because PV+ interneuron activation during retrieval did not impair OLM performance, we tested whether this manipulation affected retrieval-associated hippocampal network activity. C21 was administered to transduced *PV-CRE* mice 30 min prior to testing, and all animals were sacrificed 90 min after testing to measure hippocampal cFos expression patterns associated with memory retrieval. hM3Dq mice showed a higher proportion of AAV-labeled PV+ interneurons in DG, confirming effects of chemogenetic activation (Figure 8A-D). This activation increased expression of cFos among neurons surrounded with PNNs, but did not affect the overall proportion of AAV-labeled PV+ interneurons surrounded by PNNs (Figure 8E-G). While chemogenetic activation tended to decrease DG granule cell activation, particularly in the inferior blade, this effect was not statistically significant (Figure 8H-J). PV+ interneuron activation also had no significant effects on overall network activity in CA1 (Figure 9A-C) or in CA3 (although there was a trend toward increased overall cFos expression in CA3 with chemogenetic manipulation; Figure 9D-F). Taken together, these results suggest that, unlike the more pronounced effects of SST+ interneuron activation, the PV+ population in DG has little or no impact on dorsal hippocampal network activity during spatial memory encoding and retrieval.

**Figure 8:**
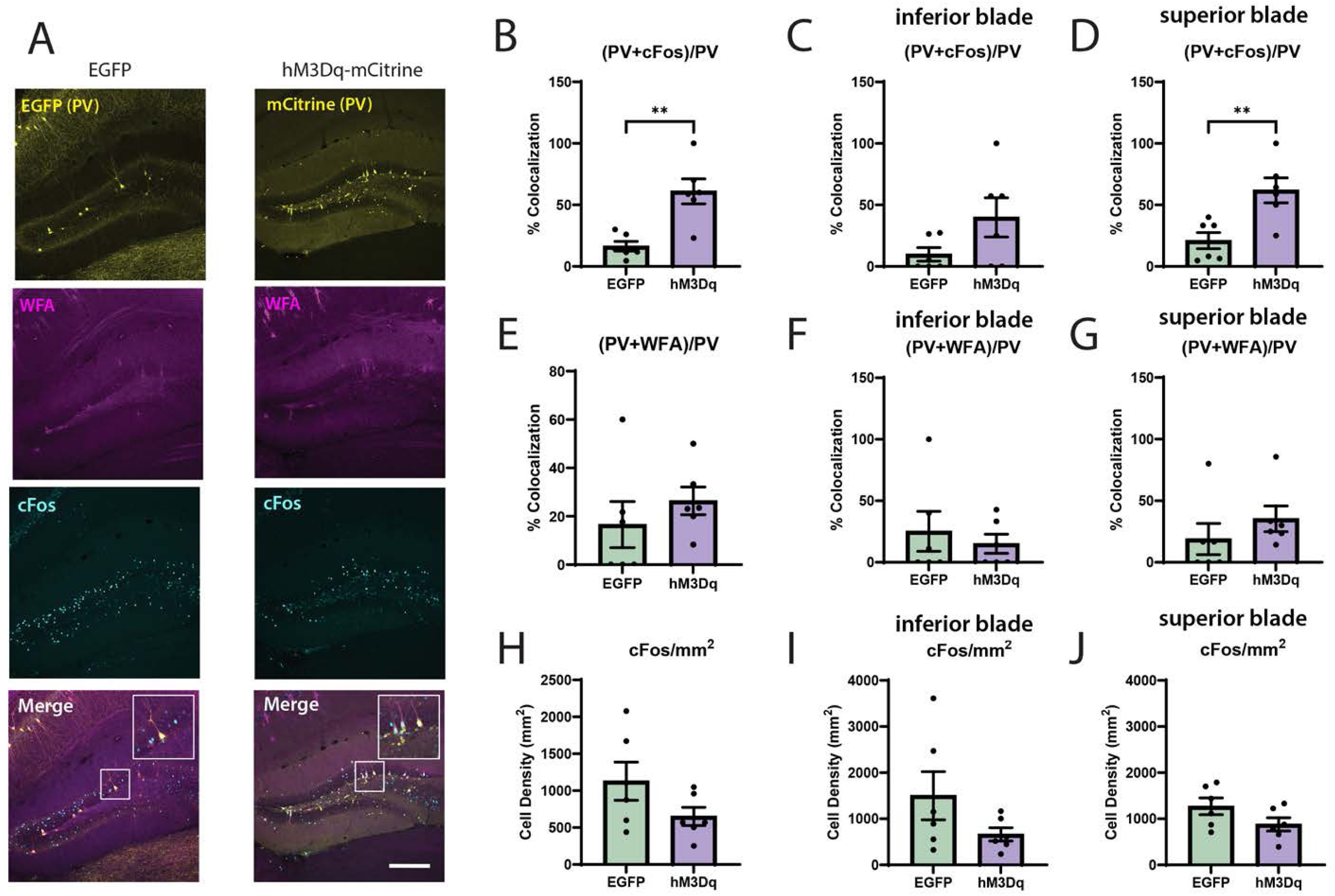
DG PV+ interneuron activation during OLM retrieval does not affect overall DG network activity. (A) Representative images of DG following OLM retrieval, from EGFP- or hM3Dq-mCitrine expressing *PV-CRE* mice, showing virus-labeled PV+ interneurons, and WFA (magenta) and cFos (cyan) labeling was used as a marker for neuronal activity. (B) Chemogenetic activation increased cFos expression among PV+ interneurons (p = 0.002), which was apparent in both the inferior blade (C; p = 0.104), and superior blade of the DG (D; p = 0.0071, 2 way ANOVA, AAV×blade interaction: p = 0.457; main effect of AAV: p = 0.020; main effect of blade: p = 0.039. (E) While PV activation during OLM retrieval did not affect the proportion of WFA expressing PV cells overall (p = 0.399), it reduced perineuronal net association in the inferior blade (F; p = 0.587), but increased it in the superior blade of the DG (G; p = 0.3405), as indicated by a significant interaction effect. 2 way ANOVA, AAV×blade interaction: p = 0.023; main effect of AAV: p = 0.852; main effect of blade: p = 0.189. (H) Whereas activation did not alter the total density of cFos-positive cells in the DG granule cell layers (p = 0.1341), it might affect both blades in an opposite direction; decreasing cell activation in the inferior blade but increasing cFos expression in the superior blade of the DG (I; p = 0.152, J; p = 0.122). 2 way ANOVA, AAV×blade interaction: p = 0.247; main effect of AAV: p = 0.131; main effect of blade: p = 0.970. Scale bar = 250 µm.

**Figure 9:**
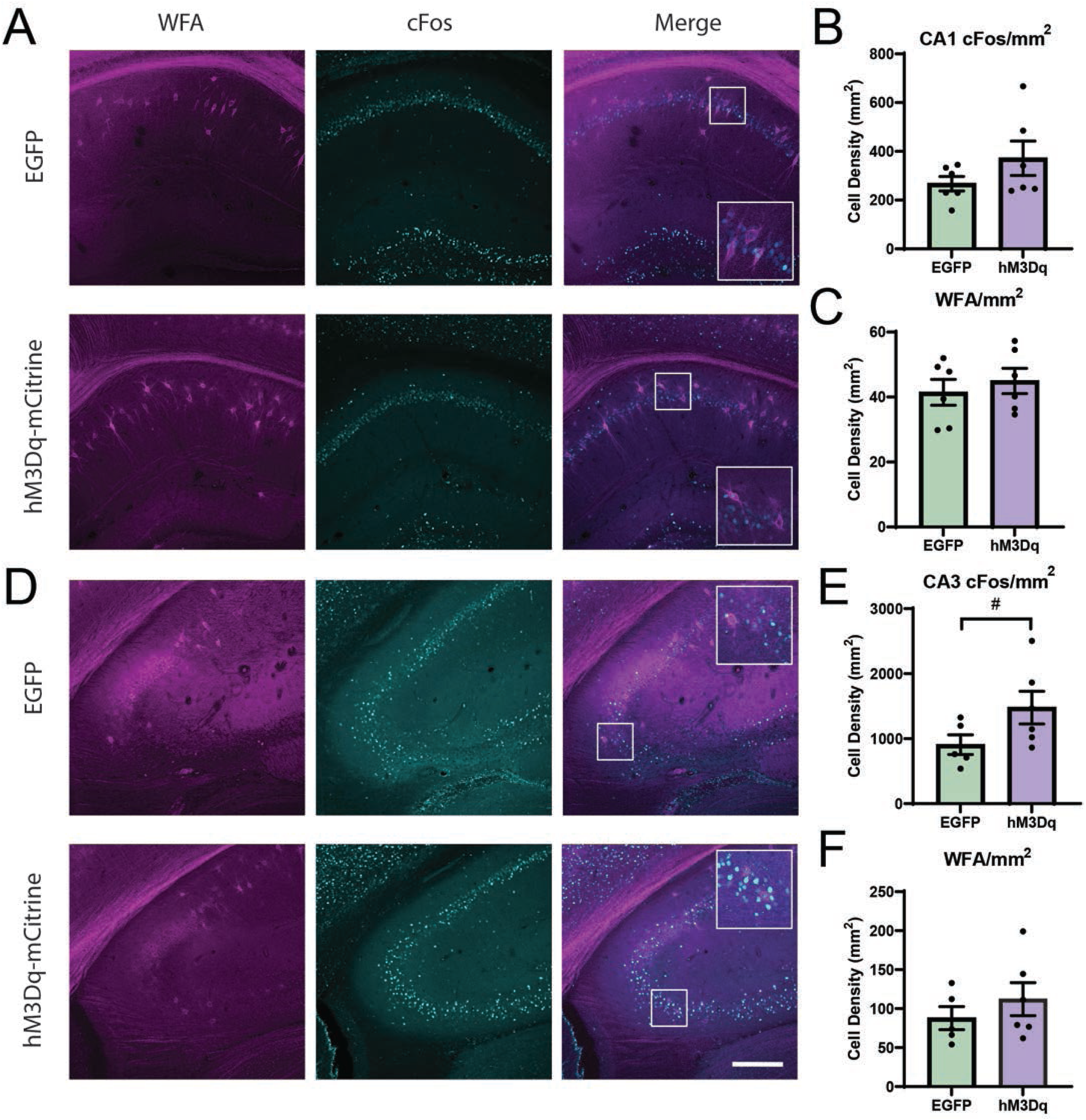
PV+ interneuron activation during OLM retrieval has no effect on CA1 or CA3 activity. (A) Representative images of CA1 region in the two groups, immunolabeled for WFA (magenta) and cFos (cyan), following OLM retrieval. Scale bar = 250 µm. There were no differences across groups in the density of cFos-positive cells (B; p = 0.203) or the density of WFA labeled cells (C; p = 0.542) in any CA1 subregions (data shown are aggregated across cell layers). (D) Representative images of CA3 following retrieval. Scale bar = 250 µm. (E) In CA3, there was a trend towards increased density of cFos-positive cells following retrieval with chemogenetic activation of PV+ interneurons (^#^p = 0.098, Student’s t-test). (F) The density of CA3 cells labeled with WFA did not differ between groups following retrieval (p = 0.391).

## Discussion

Our data show that DG SST+ interneuron activation impairs both OLM encoding and retrieval processes, while PV+ interneuron activation impairs OLM encoding alone. Somewhat surprisingly, activating neither population immediately *following* encoding had a significant negative impact on spatial memory consolidation. We find that SST+ interneuron activation suppresses activity among DG granule cells - particularly those in the inferior blade - during both phases of OLM processing. In contrast, downstream effects of interneuron activation on network activity patterns in CA1 and CA3 differ significantly when inhibition is differentially targeted to encoding vs. retrieval. For example, suppression of total CA1 cFos (and presumably, suppression of overall network activity) occurred with SST+ interneuron activation during encoding, but not with activation during retrieval. In CA3, the consequences of DG interneuron activation were more subtle, with trending effects that differed between males and females, and were again present only during encoding, and not during retrieval. These data suggest that: 1) DG network activity plays a critical role in both the encoding and retrieval of spatial memories (although not in their consolidation), 2) inferior blade DG granule cell activity may be particularly important to both processes, and 3) activation of CA1 neurons by (direct or indirect) input from DG is likely important for spatial memory encoding, while the role of this pathway in OLM retrieval may be less important. One might infer from the latter point that the processes of OLM encoding and retrieval involve distinct hippocampal neural circuits downstream of DG. This seems to be the case for other forms of memory such as contextual fear memory (CFM), where distinct neuronal populations in CA1 (which project to different output structures) are critical for memory encoding vs. retrieval ^31^. We find that hilar SST+ interneurons differentially affect the downstream activity of CA1 circuitry in the context of spatial memory encoding vs. retrieval.

One question raised by our present findings is whether, within DG, inhibitory gating of inferior blade granule cells is a rate-limiting factor for memory processing. Inhibition of granule cells suppresses encoding of both CFM ^13^ and trace eyeblink conditioning ^32^. On the other hand, the effects of granule cell inhibition during memory retrieval remain controversial ^33^. Our present data suggest that SST+ interneuron-gated DG circuits - particularly inferior blade granule cells - mediate both encoding and retrieval of spatial memory. We find that SST+ interneuron activation dramatically suppresses cFos expression in DG inferior blade during both encoding and retrieval of OLM, and that the success of OLM retrieval is predicted by the extent of inferior blade activation during OLM testing. This suggests that information processing in the inferior blade could be important for spatial memory encoding and recall. In support of this idea, recent studies from our lab have shown that successful CFM retrieval is associated with greater DG granule cell activation ^28^, and that CFM encoding causes long-lasting changes in granule cell excitatory-inhibitory balance ^19^. Moreover, sleep disruption - which interferes with all stages of hippocampus-dependent memory processing ^23^ - appears to selectively suppress inferior blade granule cells’ activity, alter their synaptic plasticity-related gene expression, and disrupt their synaptic architecture ^19,34^.

DG PV+ interneuron activation also disrupts OLM encoding. However, unlike SST+ interneuron activation, it has no impact on memory retrieval, no significant effects on cFos expression in surrounding DG granule cells, and no effect on cFos expression patterns downstream in CA1 or CA3. Together, these findings provide further evidence of the differential roles of SST+ and PV+ interneuron function within the hippocampal circuit in the context of spatial memory processing, with distinct, but functionally important, roles in memory encoding. Consistent with this functional distinction, recent work has shown that DG PV+ interneurons, unlike SST+ interneurons, do not mediate basal inhibitory signalling to DG granule cells. Rather, this interneuron population selectively provides long-lasting feedback inhibition to a subpopulation of principal neurons, semilunar granule cells ^35^, which are activated selectively during memory encoding ^36,37^. Moreover, recent data have shown that PV+ and SST+ populations in DG and CA3 differentially gate excitatory inputs to CA1 - with the SST+ population gating input from the entorhinal cortex, and the PV+ population gating input from CA3 ^38^. These mechanistic differences between DG PV+ and SST+ interneuron populations likely explain the phenotypic differences when the two interneuron types are activated. The lack of changes to hippocampal network activity with PV+ interneuron activation is likely due to the fact that within DG, this population mediates feedback inhibition only, and only to the sparse population of semilunar granule cells. In addition, if EC inputs to CA1 are only modestly affected by PV+ interneuron activation during OLM encoding, learning-associated CA1 network activity may remain intact. Intriguingly, because PV+ interneuron activation affects OLM encoding and *not* retrieval, this suggests that the neural pathways strongly modulated by DG PV+ interneurons - i.e., semilunar granule cell activity and CA3 input to CA1 - are essential only during memory encoding.

One surprising experimental outcome from these studies is the lack of OLM consolidation deficits observed when PV+ and SST+ interneuron populations are activated *following* encoding. Prior studies from our lab have shown that both PV+ and SST+ populations in the dorsal hippocampus are important for regulating CFM consolidation. For example, inhibition of CA1 PV+ interneurons in the hours following fear conditioning disrupts CFM, while rhythmic activation of this population is sufficient to rescue CFM from consolidation deficits caused by sleep deprivation ^15,16^. On the other hand, selective activation of hippocampal SST+ interneurons, which occurs in the context of sleep deprivation, disrupts consolidation of CFM, while chemogenetic suppression of SST+ interneuron activity promotes consolidation ^18^. Thus, it is somewhat surprising that activation of neither DG PV+ nor SST+ interneurons appears to affect OLM consolidation. One potential explanation for the discrepancy is the use of a purely spatial memory paradigm in our study, which unlike CFM, likely does not rely on amygdalar pathways encoding emotional salience^39^. Another possibility is that PV+ and SST+ interneuron populations in CA1 play a more important role in both OLM and CFM consolidation than those in the DG. Data from our lab and others suggest that these CA1 populations undergo significant gene expression changes and plasticity during memory consolidation, as well as during interventions known to disrupt both OLM and CFM consolidation (e.g., sleep deprivation) ^1,7,18,40,41^. Interestingly, our present data suggest that the DG network may be non-essential for “offline” mnemonic processing (i.e., consolidation), but critical for the “online” encoding and retrieval of memories. Future studies will be needed to determine whether this general principle applies to all types of hippocampus-dependent memory.

In summary, our data reveal distinct roles of DG network regulation by SST+ and PV+ interneurons in the different stages of spatial memory processing. Suppression of DG output by SST+ interneurons impairs memory encoding and retrieval, and these behavioral consequences are associated with distinct alterations to network activity downstream of DG, in CA1. PV+ interneurons’ activation within the DG impairs encoding only, likely by modulating the activity of sparse granule cell subpopulations in DG and excitatory input to other parts of the hippocampal circuit. These findings provide a roadmap for understanding how inhibitory circuits in the DG, and the DG itself, contribute to the various stages of memory processing within the hippocampus. The details of these mechanisms will have implications for understanding and treating disorders such as dementia, schizophrenia, and bipolar disorder - all of which feature neuropathological changes to these interneuron populations.

## Acknowledgments

Schematics in Figure 1A-B were created using Biorender. These studies were supported by the NIH R01NS118440, R01MH135565, a Chan Zuckerberg Initiative Collaborative Pairs grant, and a UM Biosciences Initiative grant to SJA, and awards to FR from the Michigan Alzheimer’s Disease Research Center and the Sleep Research Society Foundation.

---

**Supplementary Figure 1: Objects used for OLM.** To assess the stage-specific effects of selective SST+ or PV+ interneuron activation on OLM encoding, consolidation, and retrieval, we conducted four distinct OLM trials—each featuring unique objects and placement patterns—to isolate and examine each phase separately.

